# How non-veridical perception drives actions in healthy humans - Evidence from Synaesthesia

**DOI:** 10.1101/453134

**Authors:** Marie Luise Schreiter, Witold X. Chmielewski, Jamie Ward, Christian Beste

**Author notes:** Corresponding author Cognitive Neurophysiology, Department of Child and Adolescent Psychiatry, Faculty of Medicine of the TU Dresden, Germany Schubertstrasse 42, D-01309 Dresden, Germany. contributed equally (Marie Luise Schreiter, Witold X. Chmielewski).

## Abstract

We continually perform actions driven by our perception and it is commonly held that only objectively perceived changes within the ‘real’ world affect behaviour. Exceptions are usually only made for clinical conditions associated with hallucinations, where objectively non-existent percepts can influence behavior. Using synaesthesia as a model condition, we show that even in healthy populations irrelevant non-veridical precepts exert an effect on action. By *non-veridical* we refer to stimulus dimensions that are only subjectively perceived to be there. Applying electrophysiological (EEG) methods, we show that although these examined peculiarities are perceptual in nature, not primarily perceptual processes underlie the effects of irrelevant non-veridical perceptions on actions. Rather, high-order processes linking perceptions and motor control in medial frontal cortices reflect the underlying mechanism how irrelevant non-veridical perceptions modulate behaviour. Our results challenge assumptions about the determinants of healthy human behaviour but can be embedded within existing frameworks detailing perception action interactions.

## Introduction

We continually perform actions that are driven by our perceptions and it is commonly held that only objective changes of perceptual aspects in the ‘real’ world (that are *veridical*) can affect our behaviour. This natural understanding has dominated research to such an extent that it seems impossible that perceptual aspects lacking objective perceptual presence (that are *non-veridical*) affect our behavioural control. Exceptions are usually only made for instances of mental illnesses, associated with delusions and hallucinations. In the current study, we challenge this view by showing how actions and cognitive control are modulated by objectively non-existent but still non-pathological perceptions.

Within the field of cognitive psychology and neuroscience, the link between perception and action has been studied for decades. It is well established that perception and action are intimately intertwined and strong theoretical frameworks have been put forward focussing on this link. One of these is the theory of event coding (TEC) (Hommel et al., 2001) which states that to-be-produced events (i.e. actions) and perceived external events (i.e. stimuli) are coded by their constituting feature codes within a common format – the ‘event file’ (Hommel, 2009). Stimuli like letters, for example, may be coded by their shape (e.g. italics or bold), their colour, and their identity (e.g. A, B etc.) and all of these features are closely bound to another (i.e. integrated) to achieve a coherent perception. Responses or actions (to-be-produced events) are similarly represented by features e.g. detailing with hand or finger has to be used to execute an action. Event files establish bindings between features specifying a stimulus and features specifying an action (Hommel, 2005). The activation of an event file follows a pattern-completion logic meaning that the entire event file can be (re-) activated once a single feature of either a stimulus or a response is (re-)encountered. TEC proposes that whenever there is a perception of a stimulus that has previously been integrated in an event file, this can affect actions (Hommel, 2005). For this to happen, *all* perceived and integrated stimulus features are relevant to consider (Hommel, 2005). Notably, stimulus representations can be enriched by idiosyncratic, object-related associations (Hommel, 2009) and stimulus features can get automatically bound to a response and affect it even if its presence is neither necessary, nor useful for an action outcome (Hommel, 2005). It is therefore conceivable that actions are modulated by non-pathological, but still objectively non-existent, idiosyncratic enriched perceptions; i.e. by perceptual processes lacking perceptual presence or real-world perceptual veridicality if they are automatically activated and integrated into an event file.

A prime example of perceptual idiosyncrasy in healthy humans is synaesthesia, a condition in which the experience of a veridical percept (i.e. the inducer) consistently and automatically elicits a vivid experience in another modality (i.e. the concurrent) (Grossenbacher and Lovelace, 2001; Hubbard and Ramachandran, 2005; Rothen et al., 2018; Simner and Hubbard, 2013; Ward, 2013). Importantly, individuals with synaesthesia know that the inducer is objectively ‘there’, while the perceived synaesthetic concurrent is lacking objective perceptual presence (i.e. is *non-veridical* or illusory) (Seth, 2014). Thereby, synaesthesia can clearly be differentiated from hallucinations and delusions associated with psychopathology, and can serve as a useful tool for studying the underlying neurophysiology of healthy non-veridical perceptions relevant to the higher-order cognitive control of action (Sagiv and Frith 2013; Bouvet et al. 2017). Using synaesthesia as a model condition, we show how non-pathological perceptual feature dimensions that are not objectively perceived as present in the real world (i.e. are *non-veridical*) <u>and that are irrelevant</u> for an action nevertheless modulate the control of an action. Studies within the field of synaesthesia research have shown that an additional idiosyncratic experience triggered by an inducer can have advantageous effects on cognition, for example in the domain of memory performance (Rothen et al., 2013a, 2012). However, it has also been shown to have adverse effects on syneasthetes’ performance, for example when a presented stimulus is incongruently colored to their perceived non-veridical synaesthetic colour; comparable to a Stroop-like conflict (i.e. ‘synaesthetic Stroop-effect’) (Meier and Rothen, 2009). Notably, in these instances, colour acts as task-relevant stimulus dimension and thus, incongruently presented colors directly interfere with task-performance (i.e. colour naming).

Within the field of action selection, it has been suggested that processes operating on action-irrelevant perceptual information reflect automated processes, while processes operating on task or action-relevant perceptual information are more controlled (Botvinick et al., 2001; Ulrich et al., 2015). If non-veridical sensory content is task-irrelevant but should nevertheless affect action control, it is hypothesized that non-veridical perceptual information modulates action selection only in situations where automated processes govern action selection. No effects of non-veridical task-irrelevant perceptions should be evident under more controlled action selection processes. To examine this, we employ a novel experimental paradigm combining a Simon task with a Go/NoGo task specifically designed to measure inhibitory control performance within the context of automatic and controlled action selection modes (Chmielewski et al., 2018; Chmielewski and Beste, 2017) (refer methods section for details). Participants are asked to respond to letter stimuli (the letters ‘A’ left key and ‘E’ right key: Go) and withhold responses to the same letters printed in bold-italic style (***‘A’*** and ‘***E***’: NoGo). Thus, each letter stimulus is bound to a clear response. It has been shown that in spatially *corresponding* conditions where stimulus-action binding is mediated via automated processes (De Jong et al., 1994) response inhibition is worse, compared to spatially *non-corresponding* task conditions where stimulus-action binding is mediated via automated processes (Chmielewski et al., 2018; Chmielewski and Beste, 2017). These stimulus-response spatial congruity paradigms are referred to as Simon tasks. The current task can be completed by simply attending to the target letters’ identity (‘A’ or ‘E’) or shape (***‘A’*** and ‘***E***’) without taking into account the colour in which the letter is presented. However, for synaesthetes, processing of a letter is intimately bound to the experience (i.e. the activations) of a colour dimension. Crucially, a specific semantic representation of a letter stimulus (e.g. ‘A’) will evoke the same, consistent and automatic experience or colour (e.g. red) independent of its appearance or ‘shape’ (i.e. ‘A’, ‘a’, ‘ɑ’, ‘*A’* or ‘***A’ →*** red) (Dixon et al., 2006; Grossenbacher and Lovelace, 2001; Nikolić et al., 2011; Root et al., 2018). Importantly, even though this additionally activated synesthetic experience is task-irrelevant it is likely to be an integral part in event files in synesthetes. Within the group of synaesthetes, the idiosyncratically perceived *non-veridical* colour feature consistently accompanying the perception of a letter, will in fact, serve as an overlapping feature between different stimulus-response mappings (or ‘event files). That is, a subjectively perceived colour (i.e. red for ‘A’) represents a feature that is integrated in to two event files coding for *opposing* stimulus-response representations (‘A’: Go / ‘***A’:*** NoGo). According to TEC, feature overlap between event codes impairs behavioural control, because the cognitive representation of the stimulus-response relationship (consisting of several feature codes) has to be continuously updated and re-constructed according to the intended action (react or withhold reaction) making the behavioural response more prone to error. Anecdotal evidence suggests that synaesthetes experience a feeling of discomfort whenever they are presented with incongruence between a certain grapheme and a presented colour. Thus, by precisely modifying the presented colour of the letter, we can adapt the target stimuli to be either matching or mismatching with the subjective non-veridical colour experience of the synaesthete. This causes ‘idiosyncratic conflict’ to arise between the subjective experience of a *non-veridical* colour of the individual synaesthete and the colour of the objectively-present (veridical) letter stimulus. We predict that the presentation of a mismatching colour of the target letter will negatively affect response inhibition (i.e. it will increase false alarm rates in synaesthetes). This will only be the case in automated but not controlled action selection modes. Specifically, we predict that within the automatic condition false alarm rate should increase, while response inhibition should improve (i.e. false alarm rate decreased) within the controlled task context.

To examine whether effects of non-veridical experience specific affect the response selection level we examine neurophysiological (EEG) data. Response selection, conflict monitoring and adaptation processes have consistently been shown to be reflected by the N2 event-related potential (Folstein and Van Petten, 2008) reflecting processes of the anterior cingulate cortex and more rostral regions including the supplementary motor area (Folstein and Van Petten, 2008). The medial frontal cortex has been shown to orchestrate the connection of perception and action (Cavanagh and Frank, 2014) due to its hub-like structural and functional connection to sensory and motor areas (Cavanagh and Frank, 2014). If non-veridical experience affects cognitive control, especially these processes are hypothesized to be differentially modulated between synesthetes and controls. In synaesthetes we expect stronger N2 amplitudes in response to the existence of a non-veridical conflict between stimulus dimensions (i.e. mismatching non-veridical feature condition). By contrast, these differences in modulations are not expected for the controls. If the effects are specific for response control, there should be no effects in correlates of salience based bottom-up perceptual and attentional selection processes (i.e. P1 and N1 ERP-components) (Herrmann and Knight, 2001).

## Materials and Methods

### Participants

We included N=22 grapheme-colour Synaesthetes (19 female/ 3 male) between age 19 and 43 (31 ± 1.84 years of age) and N=22 gender-, education‐ and age - matched controls (34 ± 2.52 years of age) into the study. Matched controls were selected from the general population and screened with a questionnaire to ensure they did not experience grapheme-colour synaesthesia. We did not control for the co-occurrence of potential other types of synaesthesia (such as Sequence-Space Synaesthesia) in our participants, because the current task required responses to single letter graphemes rather than sequences of graphemes or words.

All participants had normal or corrected-to-normal vision and were screened on personal health background to ensure the sample was free of individuals previously diagnosed with any psychiatric disorders or taking regular medication (except birth control). All participants gave written informed consent prior to taking part in the experiment and received payment (12.50 £/h) or equivalent SONA credits (for Psychology students at the University of Sussex) after completion of the study. Three participants had to be excluded due to health reasons (epilepsy), insufficient synaesthetic consistency or low EEG data quality, resulting in a total of N=19 participants in each group included in the data analyses. Synaesthetes were recruited from an existing synaesthesia data base at the University of Sussex. Prior to completing the experimental task, synaesthetic consistency was checked using the Eagleman Battery (Eagleman et al., 2007; Rothen et al., 2013b). The Eagleman Battery is a set of online tests (freely accessible on http://www.synesthete.org) to determine the genuineness of different types of synaesthesia. Synaesthetes are asked to indicate the colour concurrent they perceive for graphemes 0-10 and A – Z on a colour picker resulting in an RGB-codes for each grapheme which consistency is calculated. The grapheme-colour consistency of our sample of synaesthetes was confirmed by an average score of 0.74 ± 0.12, which was significantly below the proposed cut-off value of 1.43 (Rothen et al., 2013b) (t(18) = −15.13, p < .001).

### Task and Procedure

The experiments took place in a dimly lit shielded Faraday cage. Stimuli were presented on a PC running Presentation Software (neurobehavioral systems). Participants were seated in front of a twenty-two-inch cathode ray tube monitor running at 100 Hz refresh rate. Participants executed responses via the left and the right “CRTL button” on a low latency keyboard. The monitor was placed at eye-height, and viewing distance was 60 cm.

Participants were asked to place their left index finger on the left CRTL key and their right index finger on the right CRTL key. A white fixation cross was continuously presented on a dark-grey background in the centre of the screen. The fixation cross was always accompanied by white frame boxes presented on the same vertical level in a distance of 1.1° visual angle left and right of the fixation cross.

The ‘Simon-Go/NoGo’ task (Chmielewski et al., 2018; Chmielewski and Beste, 2017) requires participants to respond to single target letter stimuli presented in normal font (i.e. ‘A’ or ‘E’; Go trials), and to withhold responses when target letters are presented in bold-italics (**‘*A*’** or ‘***E’***; Nogo trials). Whenever an ‘A’ was presented, a left-hand response was required. Whenever an ‘E’ was displayed, a right-hand response was required. These responses were required, irrespective of the spatial position of the letter stimulus, i.e. irrespective of whether the letter appeared in the left or right white frame box on the screen. Thus, the only task-relevant stimulus features for completing the task were the letters’ identity (‘A’ or ‘E) and shape (bold-italics style) and all participants were instructed that these are the only relevant feature dimensions to attend to. The task consisted of the following conditions; a spatially *corresponding* condition in which the letter-stimuli were presented on the same side of the hand carrying out the response, and a spatially *non-corresponding* condition in which the stimuli were presented on the opposite side to the hand carrying out the response. This variation creates the Simon component of the task, which applies to both, Go and NoGo trials. Essentially, Simon conflicts (i.e. spatial non-correspondence) have been shown to compromise response execution performance (i.e. hits on Go trials), but to improve inhibitory control performance (i.e. restraint on NoGo trials) (Chmielewski et al., 2018; Chmielewski and Beste, 2017).

In order to examine the effects of non-veridical perceptual features (i.e. the synaesthete-specific subjective perception of a specific colour (concurrent) when perceiving a certain letter (inducer)), this paradigm was specifically adapted for each synaesthete by adapting the colour in which the stimuli (‘A’ or ‘E’ - Go trials, and ‘*A’* or ‘*E’* - NoGo trials) were displayed (refer Figure 1 for details).

**Figure 1.**
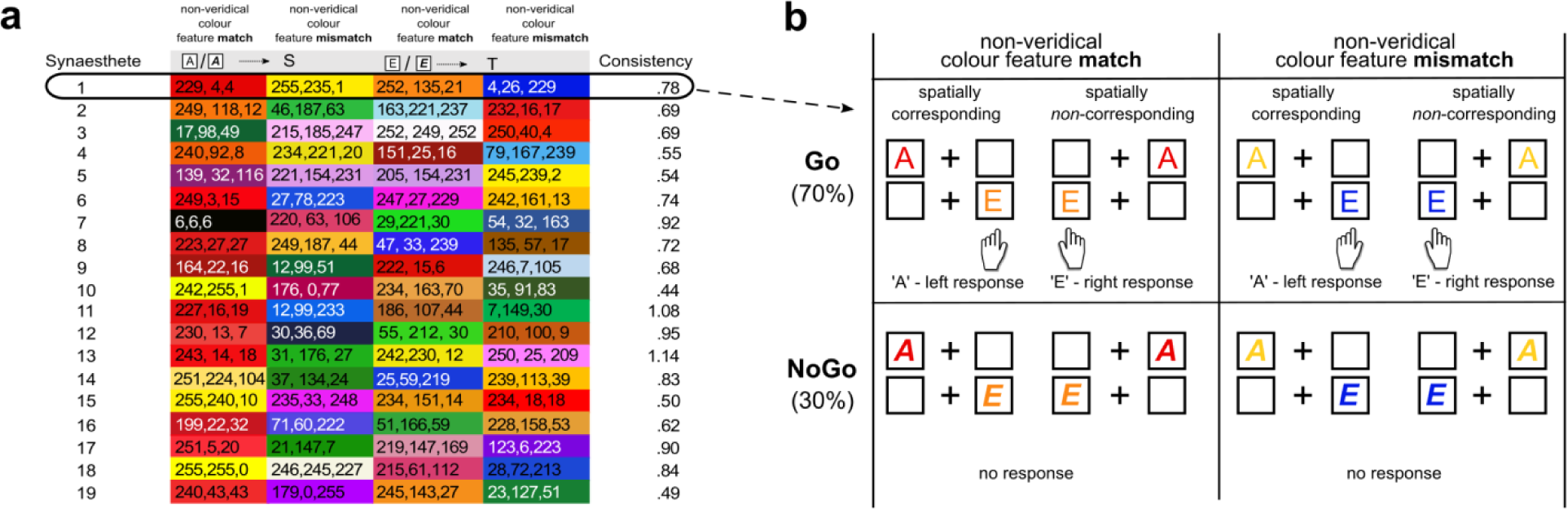
Schematic illustration of the experimental procedure. (a) Depiction of the stimulus colour used for each individual synaesthete (N=19) and their matched controls. Shown are the precise RGB codes of the non-veridical colours associated with the letters ‘A’,’E’ (target), ‘S’ and ‘T’ (idiosyncratic distractor colour) for each synaesthete and the corresponding grapheme-colour consistency scores as obtained from the Eagleman battery test. A colour consistency score below 1.43 indicates genuine synaesthesia. The stimuli in the Simon NoGo task were created on the basis of these individually determined colours. The stimuli used for synaesthete particpant 1 in the Simon NoGo paradigm are shown Figure part (b): In the ‘non-veridical colour feature **match** condition’, the colour of the presented target letter (‘A’/’E’,‘***A’***/***’E’***) was adjusted using the RGB code for the letters ‘A’ and ‘E’ (i.e. for synaesthete 1, red and orange). In the ‘non-veridical colour feature **mismatch** condition’, the colour of the presented stimuli (‘A’/’E’, ‘***A’***/***’E’***) was adjusted using RGB code for the target letters ‘S’ and ‘T’ (i.e. for synaesthete 1, yellow and blue) causing ‘idiosyncratic interference’ (please note, the letters ‘S’ and ‘T’ were never presented on screen). For Go-trials participants were instructed to respond to letters (‘A’/’E’) when presented in normal font by pressing the left button (for ‘A’) and the right button (for ‘E’) irrespective of the location (‘spatially corresponding/‘spatially non-corresponding). For NoGo-trials, participants were asked to withhold their responses whenever the letters were presented in ***bold-italics*** style (‘***A’/’E’***). On one half of the presented trials, the colour of the target stimulus exactly matched with the non-veridical perception of the synaesthete, on the other half the letters were presented in a mismatching colour. Colour never represented a task-relevant stimulus dimension.

For each synaesthete (and their matched control) the target letters (‘A’ or ‘E’) were either displayed in colours, which were in accordance, or in conflict with the non-veridical colour this specific synaesthete perceives when being presented with the ‘A’ or ‘E’ letter. By means of this manipulation, in conditions with *matching non-veridical colour features* the letter A’/ ‘***A***’ (’E’/ ‘***E’***) was presented in the *matching* synaesthete-specific RGB-code (as obtained from the Eagleman battery) of the non-veridical (automatically perceived) colour for the letter ‘A’/ ‘***A***’ (’E’/ ‘***E’***). For participant 1, for example, ‘A’/ ***A***‘s were presented in red, and ‘E’/ ‘***E‘s*** in orange (see figure 1). For conditions with *mismatching non-veridical colour features* the letter A’/ ‘***A***’ (’E’/ ‘***E’***) was presented in the RGB-codes of the synaesthete-specific synaesthetic colour concurrent of the letter ‘S’ (‘T’), hence creating interference between the subjectively perceived *non-veridical* colour of this specific letter and the actual colour, in which the stimulus was presented. For participant 1, for example, ‘A’/ ***A***‘s were presented in yellow instead of the synaesthetically perceived red, and ‘’E’/ ‘***E‘s*** in blue instead of orange (see figure 1). Hence, an ‘idiosyncratic conflict’ between the objectively presented colour of the target, and the idiosyncratically perceived non-veridical colour of the stimulus was specifically created for each synaesthete in the *mismatching non-veridical* conditions. This manipulation created the following eight experimental conditions: corresponding Go trials with matching or mismatching non-veridical colour features, non-corresponding Go trials with matching or mismatching non-veridical colour features and corresponding NoGo trials with matching or mismatching non-veridical colour features, non-corresponding NoGo trials with matching or mismatching non-veridical colour features. According to the TEC, specific stimulus feature codes are connected with specific response feature codes in event files (Hommel, 2009). In the context of this task, for both synaesthetic participants and controls task-relevant features are ‘identity’ of the letter (i.e. ‘A’ or ‘E’, relevant for Go-trial execution) indicating a specific response (left or right), and ‘shape’ (**‘*A’*** or ‘***E’***, relevant for NoGo trial identification) indicating whether to execute, or to withhold a response. Stimulus ‘location’ (left or right frame box) and even more importantly, the stimulus ‘colour’ were task-irrelevant features. For synaesthetes, however, the involuntary perception of an additional synaesthetic colour concurrent is intimately coupled to the ‘identity’ of the target letter (Root et al., 2018). Hence, a task-irrelevant stimulus feature, namely the idiosyncratically perceived ‘non-veridical colour’, is automatically activated in synesthetes whenever a letter is presented to them. For synaesthetes, the ‘non-veridical colour’ and is thus likely to be a fundamental part of the event file. In cases where the objective colour of the presented letter does not match the non-veridical colour coupled to the ‘identity’ of the target letter (conditions with *mismatching non-veridical colour features*), there will be an interference between the non-veridical and the veridical (objectively presented) colour feature dimension. Therefore, non-veridical stimulus features should impact on executive control. As ‘non-veridical colour’ is not perceived by controls, such interference should only apply to synaesthete participants. Furthermore, differential effects are to be expected in synaesthetic participants for Go and NoGo trials. This is based on the fact that for NoGo trials, the stimulus feature ‘identity’ essentially becomes task-irrelevant (A or E), as inhibition is required in response to the letters’ ‘shape’ (***bold-italics*** vs. normal font) disregarding its presented ‘identity’. Therefore, the ‘non-veridical colour’ should differentially affect Go and NoGo trial performance. Similarly, as the feature ‘identity’ might be less important for task performance in corresponding than in non-corresponding trials, performance differences might be expected based on the ‘location’ of the trial.

Before conducting the experiment, a standardized exercise of 40 trials using white letters was conducted to familiarize participants with the task. The experiment consisted of seven blocks of 160 trials with an equal distribution of pseudorandomized corresponding or non-corresponding, matching and mismatching non-veridical Go (70%) and NoGo (30%) trials (please refer to supplementary Tables 1 and 2 for details regarding the precise number or trials per condition and for the whole experiment). This Go/NoGo ratio was chosen to ensure that we establish the classic characteristic of a Go/NoGo-task, i.e. the induction of a pre-potent response tendency, which is based on the higher probability of Go trial in comparison to NoGo trial occurrence (Chmielewski et al., 2018; Chmielewski and Beste, 2017). Each trial began with the presentation of a letter for 200 ms. For Go trials a response was required within 800 ms or the trial was counted as a miss. In contrast to that, NoGo trials were considered as false alarms if any response was obtained within this time interval. The inter-trial-interval was jittered between 850 – 1050 ms. The experiment consisted of 1120 trials and took around 40min to complete depending on breaks taken by the participants. Participants were actively encouraged to take breaks in order to avoid fatigue.

### EEG recording and analysis

The EEG was recorded at a sampling rate of 256 Hz using a 64 ‐channel Refa 8 amplifier and 64-channel Waveguard EEG caps (both from ANT Neuro, Enschede, The Netherlands).

A band pass filter from 0.5 to 20 Hz (with a slope of 48 dB/oct each) and a notch filter of 50 Hz were applied. Following that, a raw data inspection was conducted manually to reject technical artefacts from the EEG, before an independent component analysis (ICA; Infomax algorithm) was conducted. Using ICA, horizontal and vertical eye movement, blinks and pulse artefacts were corrected in the EEG data. After these pre-processing steps, cue-locked segments were formed. Only correct trials were included in the data analysis; i.e. Go trials were included whenever a correct response was given in a time window of 800 ms after stimulus onset. Nogo trials were included when there was no response within 800 ms after stimulus onset. Segments started 500 ms prior to the locking point (cue onset was set to time point 0) and ended 1000ms thereafter, resulting in an overall segment length of 1500 ms. Afterwards, an automated artefact rejection was applied for all the segments. Activity below 0.5 μV in a 100ms period and a maximal value difference of 200 μV in 200ms within the epoch were used as rejection criteria. If an artefact was detected in a trial, the trial was discarded. To eliminate the reference potential from the data and to re-reference the data, we applied a current source density (CSD) transformation, which also serves as a spatial filter resulting in values for amplitudes in μV/m^2^. The CSD-transformation also works as a spatial filter that helps to find electrodes which best reflect neurophysiological processes during the paradigm (Kayser and Tenke, 2015). The segmented conditions were ‘non-veridical matching corresponding Go’, ‘non-veridical matching *non-*corresponding Go’, ‘non-veridical *mis*matching corresponding Go’, ‘non-veridical *mis*matching *non*-corresponding Go trials’, non-veridical matching corresponding NoGo’, non-veridical matching *non*-corresponding NoGo’, ‘non-veridical *mis*matching corresponding NoGo trials’, and ‘non-veridical *mis*matching *non*-corresponding NoGo trials’. A baseline correction from ‐200ms to 0 prior to target onset was applied on the relevant ERP components: P1 (at P7 & P8: 108– 118 ms after target presentation onset), N1 (at P7 & P8: 162 – 182 ms), N2 (at C4: 292 – 302 ms), P3 (at Pz: 511 – 517 ms) were identified by means of scalp topography. Within these intervals, the mean amplitude was calculated. This choice of electrode and time windows included in the data analysis was validated using a statistical approach outlined in Mückschel et al. (2014). Doing so, the above time intervals were taken and the mean amplitude within the defined search intervals was determined for each of the 60 electrode positions. Then, to compare each electrode against an average of all other electrodes, Bonferroni correction for multiple comparisons (critical threshold, p = .0007) was used. Only electrodes which displayed significantly larger mean amplitudes (i.e., negative for the N-potentials and positive for the P-potentials) when compared to other electrodes were chosen. This procedure revealed the same electrodes as those chosen by visual inspection.

### sLORETA

For source localization sLORETA (standardized low resolution brain electromagnetic tomography; (Pascual-Marqui, 2002) was applied. This algorithm provides a single linear solution for the inverse problem without localization bias (Marco-Pallarés et al., 2005; Sekihara et al., 2005). The validity of sources estimated via sLORETA analysis has been corroborated by evidence from fMRI and EEG/ TMS-studies (Dippel and Beste, 2015; Sekihara et al., 2005). The computation of the standardized current density at each voxel was executed using the MNI152 template (Fuchs et al., 2002). The sLORETA images (partitioned into 6239 voxels at 5 mm spatial resolution) of the intracerebral volume in adolescents were contrasted with the images from adults. This comparison was based on statistical nonparametric mapping utilizing the sLORETA – built – in voxel – wise randomization test with 2000 permutations (*p* < .01, corrected for multiple comparisons). Significant differences between voxels in contrasted conditions were located in the MNI brain (www.unizh.ch/keyinst/NewLORETA/sLORETA/sLORETA.htm).

### Statistical analysis

Behavioural data were analysed using mixed effects ANOVA for Go and NoGo trials separately including the within-subject factors ‘congruency’ (corresponding vs. non-corresponding) and ‘non-veridical colour feature match’ (synaesthetically matching colour vs. synaesthetically mismatching colour) and ‘group’ (Synaesthetes vs. Controls) as between-subject factor. For Go trials we analysed accuracy (%) and hit reaction times (ms). Likewise, for NoGo trails we analysed accuracy and hit reaction times only, as the small number of false alarms per condition did not allow for a meaningful interpretation of false alarm reaction times. For the neurophysiological data, the factors ‘condition’ (Go vs. NoGo) and ‘electrode’ were added into the model. Greenhouse-Geisser correction was applied where appropriate and post-hoc tests were Bonferroni-corrected. For all descriptive statistics, the standard error of the mean (SEM) is given as a measure of variability.

## Results

### Non-veridical stimulus features affect behavioural control during automated responses

A summary of the behavioural data, i.e. mean reaction times (ms) and accuracy (%) for the congruency conditions and all conditions is shown in Figure 2 (please, refer to supplementary Figure 1 a/b for plots of hit reaction time data of all conditions). For the Go trials, the mixed effects ANOVA on the hit rate revealed a main effect of “spatial congruency”, (F(1,36) = 10.63, p =.002, η_p_^2^ = .228). Responses were more accurate in the spatially corresponding condition (95.13 % ± 0.91) than in the spatially non-corresponding condition (93.55 % ± 0.92). There were no other significant main or interaction effects (all F ≤ 3.10; p ≥ .09). For the reaction times (RTs), we also found a significant main effect of “congruency” (F(1,36) = 17.56, p <.001, η_p_^2^ =.328), showing that RTs were significantly shorter (faster) on corresponding Go-trials (586 ms ± 14.33) compared to non-corresponding Go-trials (601 ms ± 13.74). There were no other significant main or interaction effects (all F ≤ 2.08; p ≥ .06).

**Figure 2.**
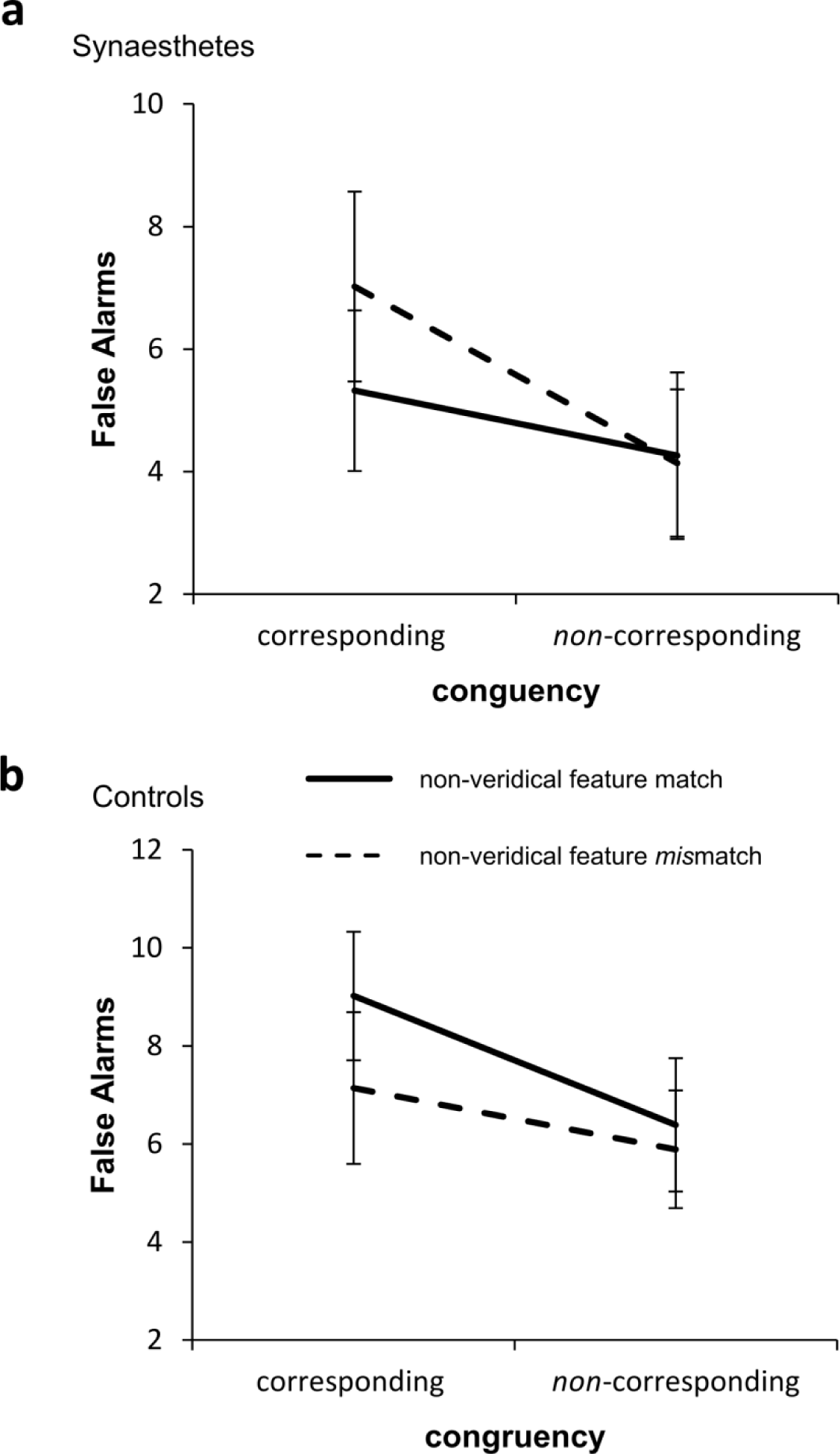
Interaction plots showing the behavioural results for both groups of participants. (a) Mean false alarm rate (in %) data for the group of synaesthetes (mean ± SEM) and (b) mean false alarm rate (in %) data for the control group (mean ± SEM). Results are shown for the spatially corresponding and spatially non-corresponding condition as well as for each non-veridical colour feature conditions (matching = solid line, mismatching = dashed lines).

However, in a Go/NoGo-task, false alarm rates reflecting inhibitory control performance represent the most important behavioural parameter. For the false alarm rates, we found a significant main effect of “spatial congruency” (F(1,36) = 11.62, p = .002, η_p_^2^ =.913) showing that the false alarm rate was higher in corresponding trials (7.12 % ± .93), compared to non-corresponding trials (5.17 % ± .85), i.e. in spatially consistent stimulus-response mappings inducing automatic response tendencies and thus, increasing false alarm rates. Importantly, there was as a significant interaction of “congruency x non-veridical feature match x group”, (F(1,36) = 8.70, p = .006, η_p_^2^ = .195). To further analyse the effects of non-veridical matching compared to non-veridical mismatching stimulus features on false alarm rates, we calculated the difference values for each group (i.e. ‘colour feature match’ minus ‘colour feature mismatch’) for the corresponding as well as the non-corresponding condition. The resulting difference values were compared using independent samples t-tests. Significant differences between the groups were revealed comparing controls (1.88 % ± 4.87) and synaesthetes (-1.70 % ± 5.42) on corresponding trials (t(36) = 2.14, p <.05). On non-corresponding trials, differences between non-veridical feature matching and mismatching trials between controls (0.50 % ± 3.87) and synaesthetes were not significant (0.13 % ± 3.68), (t(36) = 0.31, p = .761). This shows that, for synaesthetes, a stimulus inducing ‘idiosyncratic conflict’ (i.e. a stimulus with perceived mismatch between the presented and a non-veridical colour dimension on the same side as the response) acts as a distracting feature which drives up the false alarm rate specifically in this condition.

### Neurophysiological Data

#### No differential effect of non-veridical stimulus features during perceptual and attentional selection processes

The P1 and N1 ERP-components are shown in Figure 3a and 3b.

**Figure 3.**
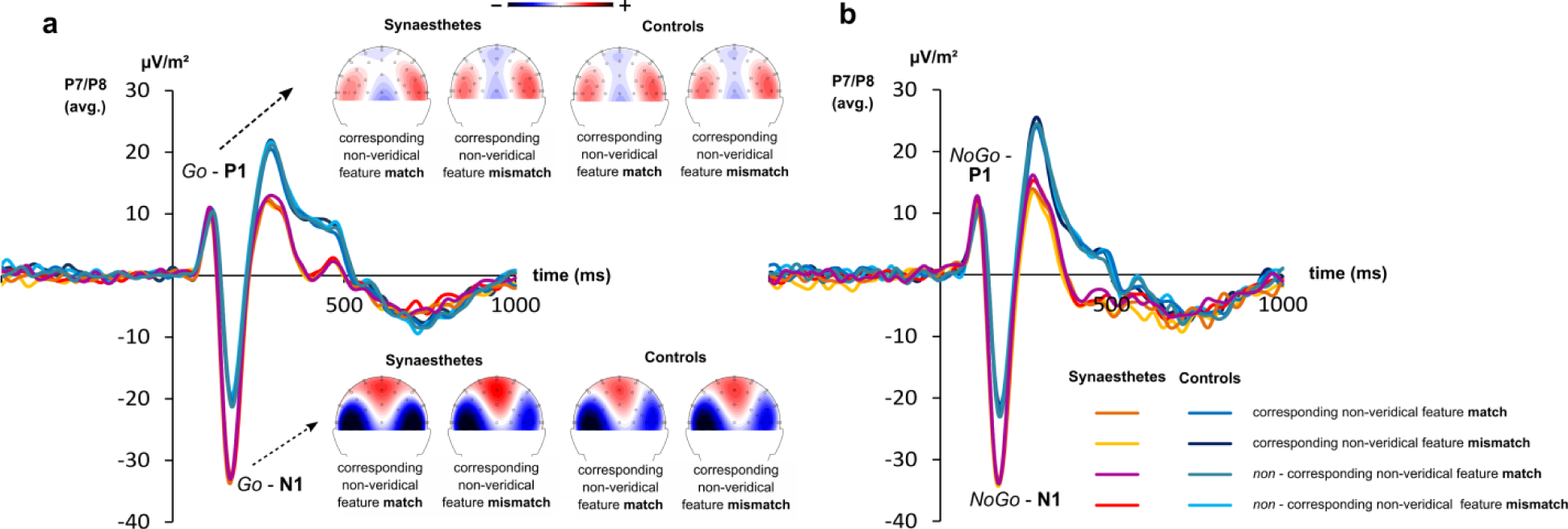
Event-related potential (ERP) P1 and N1-components averaged across electrodes P7 and P8 for each group of participants. (a) P1 and N1 on Go trials. (b) P1 and N1 on Nogo trials for all experimental conditions. Example scalp topographies are shown for non-veridical conditions since these are the most important conditions in the study. Within the topographies red colours denote positive amplitudes, blue colour negative amplitudes. The different colours of the ERP traces represent the experimental conditions for both groups. Warm colours are used to show the experimental conditions in the group of synaesthetes, cool colours are used to show the experimental conditions in the group of controls. Time point zero denotes the point of stimulus presentation.

Concerning the P1-ERP component as a correlate of perceptual gating processes (Herrmann and Knight, 2001), the analysis of the amplitude data at electrodes P7 and P8, only revealed a significant interaction effect of “electrode x spatial congruency” (F(1,36) = 43.66, p = .046, ηp = .104). To follow up this interaction, we employed independent samples t-test comparing P1 at electrode P7 on corresponding trials and electrode P8 on corresponding trials as well P7 on non-corresponding trials and electrode P8 on non-corresponding trials. We found that on corresponding trials, P1 was significantly greater (i.e. more positive) at electrode P8 (11.83 μV/m^2^ ± 1.73) compared to electrode P7 (8.77 μV/m^2^ ± 1.54), (t(36) = −2.24, p >.05). For the non-corresponding condition, expression of P1 did not significantly differ between electrode P7 (10.10 μV/m^2^ ± 1.43) and P8 (12.10 μV/m^2^ ± 1.70), (t(36) = −1.70, p =.11). Interestingly, there were no other main or interaction effects including the factor “group” (all F ≤ 4.04; p ≥ .53). This shows that within the automatic task context (i.e. spatially corresponding stimulus response mappings) the P1 was more lateralized towards the right hemisphere.

Concerning N1 ERP-component as a correlate of bottom-up attentional selection processes (Herrmann and Knight, 2001), the analysis of the amplitude data at electrodes P7 and P8, there was a significant main effect of “electrode”, (F(1,36) = 7.38, p = .047, η_p_^2^ = .105) showing that N1 amplitudes were larger (i.e. more negative at electrode P7 (-4.67 μV/m^2^ ± .50) compared to electrode P8 (-2.85 μV/m^2^ ± .59). There also was a significant main effect of “Go vs. NoGo” (F(1,36) = 8.68, p = .006, η_p_^2^ = .194) showing that N1 was more negative in NoGo condition (-27.36 μV/m^2^ ± 2.51) compared to the Go condition (-26.00 μV/m^2^ ± 2.58). We also found a significant main effect of “group” (F(1,36) = 4.95, p=.32, η_p_^2^ = .121), showing that N1 amplitude was more negative in synaesthetes (-32.32 μV/m^2^ ± 3.56) compared to controls (-21.04 μV/m^2^ ± 3.56). The source localization shows that the precuneus (BA19) revealed stronger activity in synaesthetes than controls (refer supplementary figure 2) and reflect unspecific changes in perceptual/attentional processes related to synaesthesia itself. The only significant interaction was an interaction “electrode x Go vs. NoGo x congruency” (F(1,36) = 52.02, p = .024, η_p_^2^ = .133). However, post-hoc tests revealed that this interaction only revealed that N1 amplitude differences between Go and Nogo trials were larger at electrode P8 on incongruent than congruent trials. This, however, does not explain the differential group effects observed at the behavioural level. As with the P1 data, there were no other main or interaction effects including the factor “group” (all F ≤ 0.94; p ≥ .53).

This lack of modulatory effects on the P1 and N1 by the factor “group” is corroborated by a Bayesian analysis of the data. The approach proposed by Masson et al. allows estimating the relative evidence for different statistical models from sums-of-squares data used in the ANOVA (Masson, 2011). This analysis revealed a probability for H0 hypotheses given the data (pH0|D) of p = 0.96 thus providing strong evidence for the null hypothesis according to the criteria by (Raftery, 1995); i.e. that there were no group effects.

#### Neurophysiological mechanisms of response selection are modulated by non-veridical stimulus features

The N2 ERP-component is shown in Figure 4a and b.

**Figure 4.**
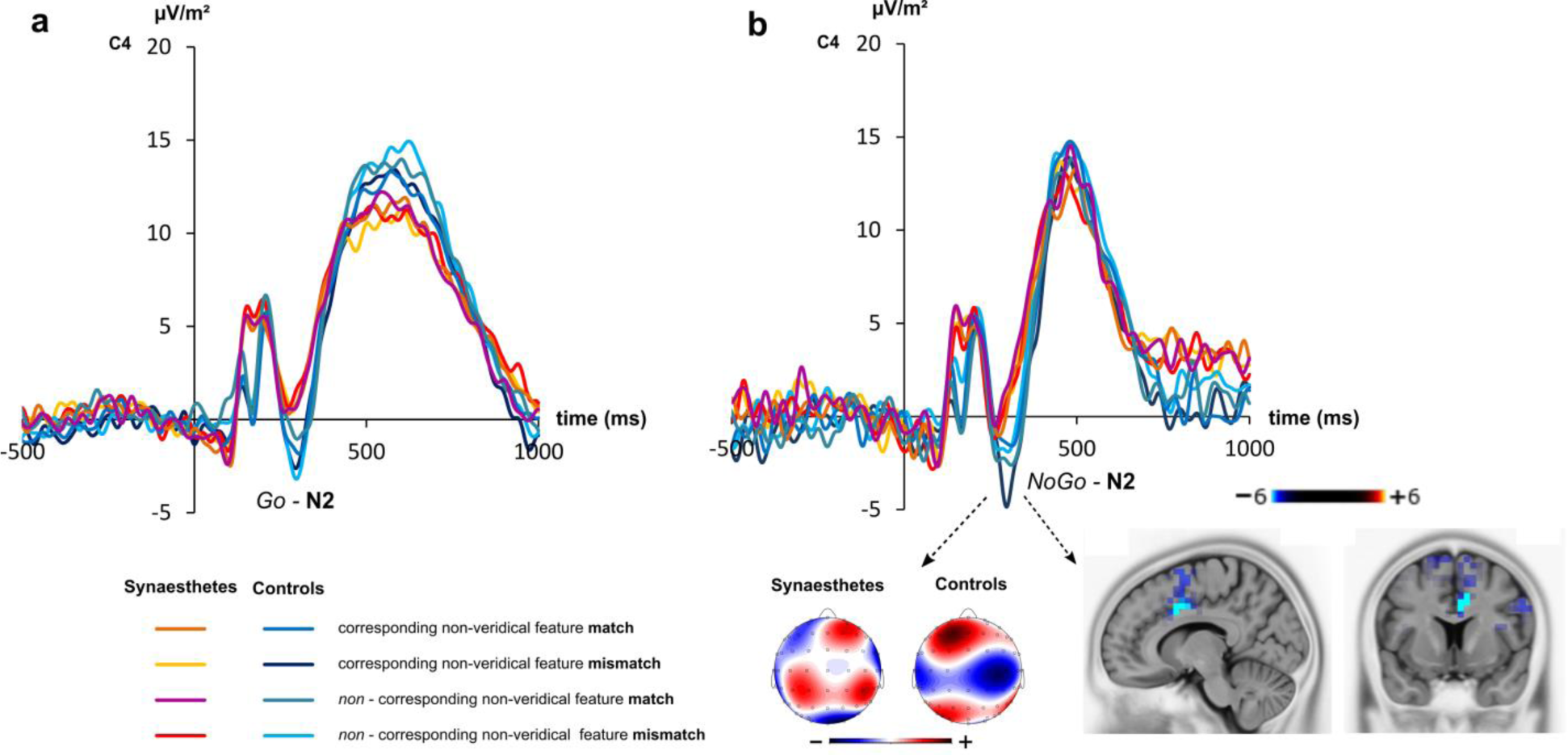
Event-related potential (ERP) N2-component at electrode C4 for both groups of participants with scalp topographies as well as a visual representation of the source localization results (sLORETA). Components are shown for (a) the Go-N2 and (b) NoGo-N2 in each experimental condition. Scalp topographies show the difference in ERP-signal between the non-veridical **matching** colour feature condition and the non-veridical **mismatching** colour feature condition. Within the topographies, red colours denote positive amplitudes, blue colours negative amplitudes in the difference. The different colours of the ERP traces represent the different experimental conditions for both groups. Warm colours are used to show the experimental conditions in the group of synaesthetes, cool colours are used to show the experimental conditions in the group of controls. Time point zero denotes the point of stimulus presentation. The sLORETA results indicate the source of the differential effect between experimental conditions and groups in the NoGo-N2 in the anterior cingulate cortex (ACC) (corrected for multiple comparisons using SnPM, p < .01). The colours denote critical t-values.

Concerning N2 ERP-component amplitudes at electrode C4 we found a significant main effect of “condition” (F(1,36) = 51.22, p < .001, η_p_^2^ = .587), showing that N2 amplitudes were higher (i.e. more negative) on NoGo-trials (-1.28 μV/m^2^ ± 1.03) compared to Go-trials (17.54 μV/m^2^ ± 2.39). Importantly, there was a significant interaction of ‘Go vs. NoGo x non-veridical feature match x congruency x group’, (F(1,36) = 4.20, p =.023, η_p_^2^ = .637). To analyse this interaction in more detail, we calculated mixed effects ANOVAs for Go‐ and the NoGo trials separately. For Go trials, there were no significant main or interaction effects (all F ≤ 0.13; p ≥ .5). For NoGo trials, there was a significant interaction “non-veridical feature match x congruency x group” (F(1,36) = 5.54, p =.024, η_p_^2^ = .133). To further analyse the effects of non-veridical matching compared to non-veridical mismatching stimulus features in NoGo trials, we calculated the difference values for each group (i.e. ‘synaesthetic feature match’ *minus* ‘synaesthetic feature non-match’) for the corresponding as well as the non-corresponding condition. The resulting difference values were compared by means of independent samples t-test. For corresponding trials, differences in N2 amplitudes at electrode differed significantly between controls (3.34 μV ± 5.64) and synaesthetes (-0.57 μV/m^2^ ± 4.12), (t(36) = 2.44, p=.02). For non-corresponding trials, differences of N2-amplitude between synaesthetes (0.67 μV/m^2^ ± 4.91) and controls (-0.53 μV/m^2^ ± 5.59) did not reach significance, (t(36) = −0.70, p = .50). The sLORETA analysis revealed that this differential effect in N2 amplitudes between controls and synesthetes was associated with activation differences in the anterior cingulate cortex. All other main effects or interaction were not significant (all F ≤ 1.14; p ≥ .4).

The P3 ERP-component at electrode Pz, as well as FCz is shown in Figure 5 a and b.

**Figure 5.**
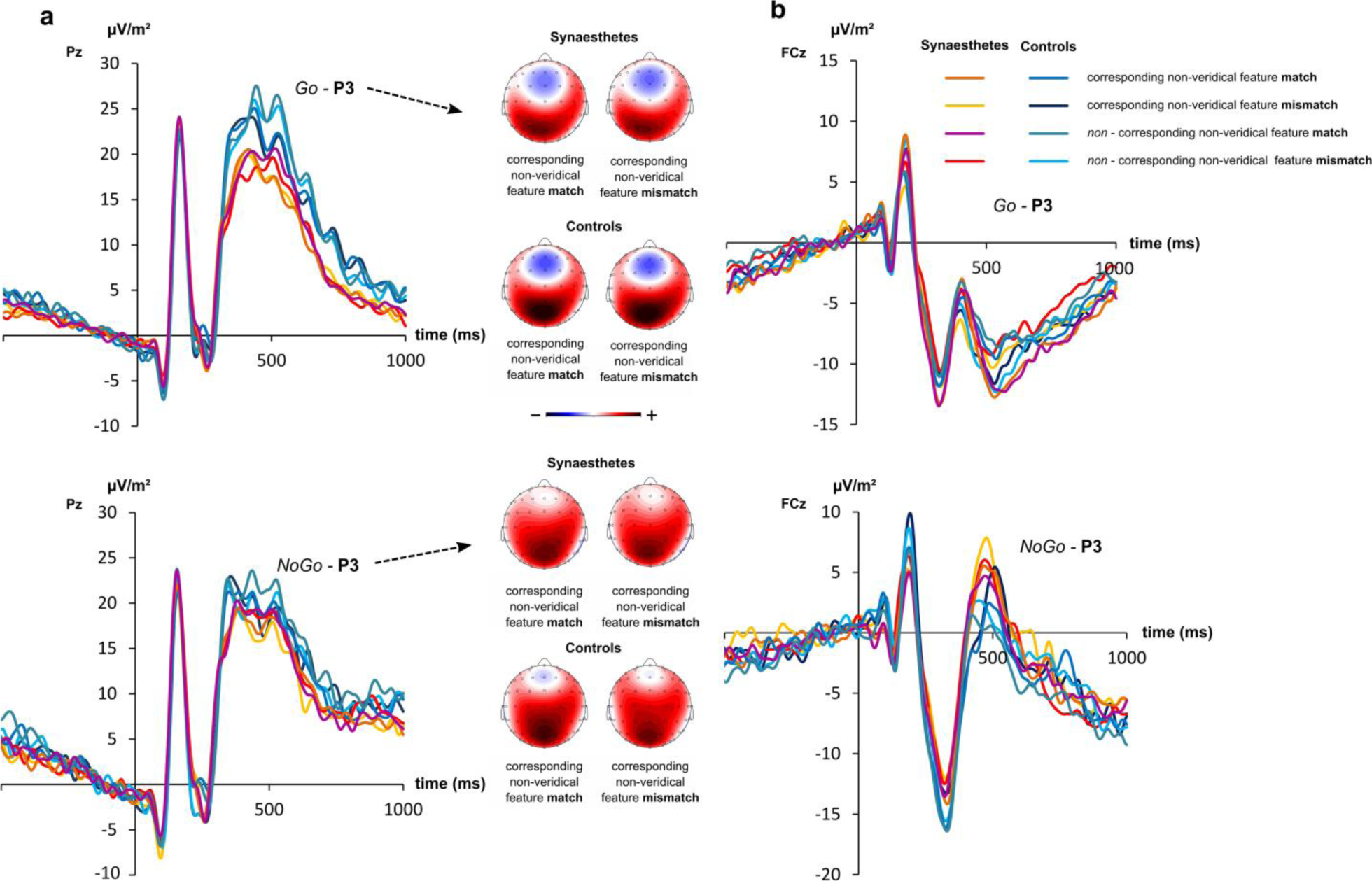
Event-related potential (ERP) P3-components at electrodes (a) Pz and (b) FCz for both groups of participants with scalp topographies. (a) Go and NoGo-P3 at C4 and (b) Go and NoGo P3 for all experimental conditions. Scalp topographies are shown to represent scalp distribution for the relevant experimental conditions. Within the topographies, red colours denote positive amplitudes, blue colours negative amplitudes in the difference. The different colours of the ERP traces represent the different experimental conditions for both groups. Warm colours are used to show the experimental conditions in the group of synaesthetes, cool colours are used to show the experimental conditions in the group of controls.

Concerning P3 at electrodes Pz and FCz we found a significant main effect of ‘electrode’, (F(1,36) = 93.81, p < .001, η_p_^2^ = .723) showing that P3 was larger at electrode Pz (20.58 μV/m^2^ ± 2.11) compared to electrode FCz (0.25 μV/m^2^ ± .39). There also was a significant main effect of ‘congruency’, (F(1,36) = 10.79, p = .002, η_p_^2^ = .231) showing that P3 was significantly smaller on spatially corresponding trials (9.80 μV/m^2^ ± 1.10) compared to spatially non-corresponding trials (11.03 μV/m^2^ ± 1.15). However, there was no interaction ‘Go vs. NoGo x non-veridical feature match x congruency x group’ (F < 0.56; p ≥ .8). Other remaining interactions are shown in the supplemental material. This lack of modulatory effects on the P3 reflecting the interaction shown for the behavioral data is corroborated by a Bayesian analysis of the data. This analysis revealed a probability for H0 hypotheses given the data (pH0|D) of p = 0.97 thus providing strong evidence for the null hypothesis according to the criteria by (Raftery, 1995).

## Discussion

In the current study, we show that actions and cognitive control processes are modulated by non-pathological perceptions that are not objectively present i.e. by experiences in which a perceptual presence or real-world perceptual veridicality is lacking. By examining synaesthesia, a perceptual phenomenon in healthy humans distinct from psychopathological delusions or hallucinations (Bouvet et al., 2017), we detail the mechanisms how this is possible. Grapheme-colour synaesthesia is a condition in which single digits, letters and words (i.e. inducer) consistently and automatically evoke an additional experience of colour (i.e. concurrent) (Grossenbacher and Lovelace, 2001; Hubbard and Ramachandran, 2005; Simner and Hubbard, 2013; Ward, 2013). This experience of a synaesthetic concurrent, despite being perceptually vivid, lacks perceptual presence; i.e. it is *non-veridical* (Seth, 2014).

Previous research has shown that experiencing an additional non-veridical percept can have adverse effects on cognition (Meier et al., 2015; Meier and Rothen, 2009). However, so far disadvantageous effects on performance were only evident in conflict tasks when the task-relevant colour stimulus was incongruent to the perceived non-veridical synaesthetic colour; and thus, directly causing interference with the task at hand (i.e. with colour categorization) (Meier et al., 2015). Notably, in our study, ‘colour’ (neither veridical, nor non-veridical) never was a task-relevant stimulus dimension.

The behavioural data clearly show that such non-veridical features affect performance even if there is no direct interference with task-relevant stimulus dimensions. We demonstrate this by employing a novel paradigm combining a Simon task with a Go/NoGo task to measure inhibitory control performance within the context of automatic (i.e. spatially corresponding stimulus response mappings) and controlled action selection modes (i.e. spatially non-corresponding stimulus-response mappings) modified to match (or mismatch) the idiosyncratic perception of each individual in the group of synaesthetes (Chmielewski et al., 2018; Chmielewski and Beste, 2017). In both groups, we found that RTs were shorter and accuracy was higher on spatially corresponding compared to spatially non-corresponding stimulus-response relations in Go trials (Hommel, 1993; Hommel et al., 2001; Wascher et al., 2001) and that erroneous responses on NoGo trials (i.e. false alarm rate) were increased on corresponding compared to non-corresponding stimulus-response relations is in line with previous findings (Chmielewski et al., 2018; Chmielewski and Beste, 2017). In syneasthetes, false alarm rates were increased whenever the veridical colour of the presented letter stimulus was not matching the subjectively perceived non-veridical colour (i.e. synaesthetic concurrent) induced by the presentation of the letter (i.e. the inducer). Thus, the mismatch between veridical and non-veridical colour features of the sensory content modulates executive control and impulsive behaviour in particular. Interestingly, these effects depend on the specific response selection mode: According to the dual process model (De Jong et al., 1994), behaviour is mediated via a ‘direct’ route in response to spatially corresponding stimulus-response relations, in which a more automatic stimulus-response translation is evident. Opposed to this, an ‘indirect’ route involving a controlled stimulus-response translation mediates behaviour on incongruent trials. The increase of false alarm rate in the corresponding condition seems to suggest that non-veridical perceptions only modulate behavioural control when response selection relies on rather automated processes. It seems that, although the perceptual presence (veridicality) of the perceived stimulus colour is what segregates the examined groups, effects of perceptual idiosyncracy only become evident in higher-order cognitive control functions. Furthermore, if different degrees of perceptual saliency (of the target letters) were to underlie the false alarm rate differences, saliency effects would likely improve the synaesthetes performance to *congruently* coloured target letters (i.e. idiosyncratically matching trails), which was not the case. Thus, early perceptual processes (effects of colour saliency) cannot be assumed to underlie the behavioural differences. This is corroborated by the neurophysiological data, showing that the P1 and N1 ERP-components, reflecting bottom-up perceptual gating and attentional selection processes (Herrmann and Knight, 2001), did not reveal interactive effects idiosyncratic non-veridical stimulus dimensions in line with the behavioural data. A Bayesian analysis of this data providing strong evidence for a lack of group effects for the P1 and N1 ERP-components further supportes this interpretation. Since the P1 and N1 ERP-component are well-known to be modulated by bottom-up processes e.g. driven stimulus saliency (Beste et al., 2012; Herrmann and Knight, 2001; Luck and Kappenman, 2013; Wascher and Beste, 2010) this finding also rules out that the stimulus manipulations could have led to differences in the saliency of the target letters. As shown in the examples presented in Figure 1, the capital letter A presented in yellow may be considered to be less salient than when it is presented in red or orange. The electrophysiological data shows that this does not confound our results.

Importantly, the interaction of “synaesthetic feature match x congruency x group” was evident for an ERP component reflecting higher-order response selection and cognitive control processes – the N2. In the context of response inhibition, the N2 is assumed to reflect pre-motor inhibition processes with the N2 being larger when these processes are fully deployed (Chmielewski and Beste, 2017; Falkenstein et al., 1999; Folstein and Van Petten, 2008; Nieuwenhuis et al., 2004). In controls, N2 amplitudes were more negative in idiosyncratically mismatching feature trials compared to idiosyncratically matching feature trials. This was not shown in the case of the syneasthetes. In fact, a mismatch between the presented colour of the letter stimulus and the non-veridical colour feature associated with the letter (i.e. concurrent) seemed to prevent pre-motor inhibition processes to be deployed as was reflected in worsened response inhibition performance (increased false alarm rates) whenever the task context was mediated via more automatic stimulus-response translations (‘direct’ route). The source localization analysis suggests that these modulations of the N2 are associated with the anterior cingulate cortex (ACC). This region, and the medial frontal cortex in general plays a major role for inhibitory control (Huster et al., 2013); and has been suggested to orchestrate the connection of perception and action (Cavanagh and Frank, 2014) due to its hub-like structural and functional connection to sensory and motor areas (Braem et al., 2017; Bush et al., 2000). Corroborating this, it has been shown that the N2 reflects a concomitant coding of stimulus and response-related aspects likely, mediating the binding between stimulus and response features (Chmielewski et al., 2018; Folstein and Van Petten, 2008). The neurophysiological data thus, suggest that altered stimulus-response binding processes underlie effects of sensory aspects lacking real-world perceptual veridicality on cognitive control. The idea of binding stimulus and response features to facilitate an action is central to the theory of event coding (TEC) detailing the links between perception and action (Hommel et al., 2001). TEC can also explain how non-veridical perceptual aspects modulate overt response control:

According to TEC, feature codes defining a percept and an action are represented in a common representational structure called “event file”. Event files establish bindings between features specifying a stimulus and features specifying an action (Hommel, 1993; Hommel et al., 2001). Importantly, the activation of an event file follows pattern completion logic meaning that the entire event file can be (re)activated once a single feature of either a stimulus or a response is (re-) encountered. According to TEC, an event file is built activating *all* stimulus and response features, no matter whether they are relevant to the task at hand or not (Hommel, 2005). Moreover, once a perceptual feature dimension has been integrated into an event file, it is likely to affect action. So, if stimulus representations are enriched by idiosyncratic, stimulus-related associations, the feature dimensions are automatically bound to a response even if their presence is neither necessary, nor useful for the desired outcome of the action (Hommel, 2009; Hommel et al., 2001). Following this assumption, we hypothesized that action should be modulated by task-irrelevant perceptual feature dimensions, even if these are *not* objectively perceived to be present in the real word, i.e. strong, automatic stimulus-related associations, lacking perceptual veridicality. In our experiment, the task-relevant feature dimensions of the target stimulus were the letters’ ‘identity’ (A or E) and its ‘shape’ (normal print or bold-italics). Importantly, the ‘colour’ dimensions never did serve as a task-relevant feature dimension. Yet, in grapheme-colour Synaesthesia the presentation of the letter A or E (i.e. the ‘identity’: task-relevant) inevitably and automatically triggers the experience of an additional non-veridical feature dimensions (i.e. ‘colour’: task-irrelevant). Crucially, the consistent and automatic colour experience (e.g. red) is intimately bound to the specific semantic representation of a letter stimulus (e.g. ‘A’) independent of its appearance or ‘shape’ (i.e. ‘A’, ‘a’, ‘ɑ’, ‘*A’* or ‘***A’ →*** red) (Grossenbacher and Lovelace, 2001; Nikolić et al., 2011). Thus, even though this additionally activated synesthetic experience is irrelevant to the desired action, it is likely to be an integral part of the event files of synesthetes. According to TEC, stimulus and response features are integrated into event files and prepared to before the task is performed, so that simply registering a stimulus should suffice to spread activation to the related response. Thus, the same non-veridical colour is activated and integrated into Go as well as NoGo - event files. Since the letter (i.e. ‘identity’) itself is not sufficient for deciding whether to execute or to withhold a response, and because the feature carrying this information (i.e. ‘shape’) does not elicit a non-veridical colour experience in grapheme-colour Synaesthesia. Therefore, Go and NoGo event files partially overlap in terms of their integrated non-veridical feature dimension for synaesthetes. The same non-veridical feature is part of two opposing event files. Such a partial overlap between opposing event files will cause interference leading to performance decline (Hommel et al., 2001). In our experiment this interference is amplified by presenting the inducer (i.e. letter) in a veridical colour incongruent with the concurrent (i.e. non-veridical colour) ultimately leading to a decline in inhibitory control performance. As a result, the synesthetic experience leads to interference between opposing event files and modules action selection.

In summary, we show, using synaesthesia as a model condition, how irrelevant and non-existent but still non-pathological perceptions; i.e. perceptual aspects lacking objective perceptual presence (i.e. that are *non-veridical*) can affect our actions. We provide insights into the conditions under which such a modulation is possible. We show that even though the examined peculiarities are perceptual in nature, not perceptual processes but later integration processes with motor response in medial frontal cortices reflect the underlying mechanism how actions are modulated by non-veridical perceptions. Although the results challenge common sense conceptions about the determinants of human action, the results can be explained by existent, influential theoretical frameworks detailing the link between perception and action.

## Acknowledgements

This work was supported a Grants from the Deutsche Forschungsgemeinschaft (DFG) SFB 940 project B8. We thank David Schwartzman for his technical support.

